# Human CYP2C9 metabolism of organophosphorus pesticides and nerve agent surrogates

**DOI:** 10.1101/2025.07.29.667567

**Authors:** Pratik Shriwas, Abigail Noonchester, Andre Revnew, Thomas R. Lane, Christopher M. Hadad, Sean Ekins, Craig A. McElroy

## Abstract

Of the Cytochrome P450 enzymes, the CYP2C9 variant is very important and is the cytochrome P450 (CYP) involved in the metabolism of several human drugs where it acts as a natural bioscavenger. Previously, CYP2C9 was demonstrated to remove sulfur from different organophosphorus (OP) pesticides, such as dimethoate, diazinon and parathion. In this study, we have tested the ability of CYP2C9 to degrade other OP compounds. We investigated the metabolism of OP compounds by CYP2C9 using LC-MS/MS approaches as well as the time-dependent inhibition of CYP2C9 by OP compounds using the previously developed pFluor50 fluorogenic assay. We found that CYP2C9 metabolizes thions (P=S) preferentially over oxons (P=O) and there were many OP compounds that inhibited CYP2C9 activity in a time-dependent manner. Additionally, we performed molecular docking based on the Protein Data Bank (PDB)crystal structure (1OG5) of the CYP2C9 receptor in order to gain atomistic insight from the *in vitro* experimental results. We observed a positive though moderate correlation between the calculated binding energy and the CYP2C9 metabolism of various OP compounds (R = 0.61) as well as the time-dependent changes in inhibition of OPs with binding energy (R = 0.59). Although the R values were modest,59). These *in vitro* data combined with analysis of additional OP derivatives could be used to develop artificial intelligence (AI)machine learning models to predict the metabolism of specific OP compounds by CYP2C9. This type of approach could be particularly relevant for the prediction of the metabolism of current and emerging chemical warfare agents.

## Introduction

Exposure to organophosphorus (OP) compounds results in both acute and chronic symptoms due to covalent inhibition of acetylcholinesterase (AChE) (Rendic and Guengerich, 2021). Deliberate OP pesticide exposure results in a very high number of suicide cases with approximately 150,000-200,000 deaths recorded every year worldwide (Blain, 2011; Eddleston, 2024), while also contributing to ∼15% of all suicidal deaths per year (Mew *et al*., 2017). Global pesticide use continues to increase with an estimated 3.7 million metric tons used per year in 2022 (*Global pesticide consumption 1990-2022*, accessed 27 July 2025) leading to as many as 25 million agricultural workers worldwide experiencing unintentional pesticide poisoning each year (Jeyaratnam, 1990) with significant health consequences such as cancer and neurotoxicity (Alavanja, Hoppin and Kamel, 2004). Additionally, the residual pesticide that remains on the crops, such as fruits and vegetables (Beyuo *et al*., 2024; Habran *et al*., 2024; Soman *et al*., 2024), leads to chronic OP exposure for a significant percentage of the population. More intentional OP compound poisoning comes from another class, chemical warfare nerve agents (CWNAs), wherein attacks on individuals or groups of people often results in death due to the increased toxicity of these compounds. The highest-profile tragedies include the use of Sarin in Syria in 2013 (Brooks *et al*., 2018), the assassination of Kim Jong Nam in Kuala Lumpur with VX in 2017 (Brunka *et al*., 2022), the use of a Novichok agent in an assassination attempt on Yulia and Sergei Skripal, resulting in the death of Dawn Sturgess in Salisbury UK in 2018, and most recently in the poisoning (2020) of Alexei Navalny in Russia (Haslam *et al*., 2022).

OP compounds can be divided into two major groups, oxons (P=O) or thions (P=S),, depending on whether the phosphorus atom is bonded to oxygen or sulfur, respectively (Bharate *et al*., 2010). Chemical warfare agents are typically oxons, while OP pesticides are delivered as thions and are then metabolized to the oxon form, thereby creating the active metabolite which inhibits the critical enzyme, AChE (Thompson, Prins and George, 2010). Currently, the treatment for OP poisoning includes the use of the anticholinergic agent, atropine, and the use of AChE reactivators, such as pralidoxime, obidoxime or HI-6, each of which has although limited efficacy (Eddleston *et al*., 2008; Kharel *et al*., 2020). However, due to poor crossing of the blood-brain barrier, these pyridinium oxime reactivators are not effective in the central nervous system. Quinone methide precursors mitigate many of the weaknesses of pyridinium oximes as reactivators of OP-inhibited AChE, while providing protection in the central nervous system; however, at present, such novel reactivators are still in development (Lovins *et al*., 2024) and so there is currently no adequate therapy which has been approved for OP poisoning.

Hepatic CYP450 enzymes play a critical role in phase I metabolism of xenobiotics including pharmaceuticals and are responsible for drug-drug interactions (Rendic and Guengerich, 2021). Recently, we showed that human liver microsomes (HLMs) are capable of degrading many OP pesticides and CWNA surrogates using targeted NMR and LC-MS/MS metabolomics (Agarwal *et al*., 2023). However, human HLMs consist of numerous enzymes including a number of CYP450 enzymes, esterases and amidases (Timoumi *et al*., 2019), so at present, it is unclear which specific enzymes are responsible for the metabolism of a given molecule.

In this study, we explored if CYP2C9 was capable of metabolizing OP pesticides and surrogates of CWNAs using LC-MS/MS metabolomics. Indeed, CYP2C9 is the most abundant member of the CYP2C family and contributes 20% of the hepatic CYP450 content in HLMs (Evans and Relling, 1999; Gillam and Hayes, 2013). It is also responsible for metabolism of 15% or approximately 100 clinical drugs (Isvoran *et al*., 2017). CYP2C9 haspreviously been shown to desulfurate thions, such as diazinon (Kappers *et al*., 2001), dimethoate (Buratti and Testai, 2007), disulfoton (Usmani *et al*., 2004), fenthion (Leoni, Buratti and Testai, 2008), malathion (Buratti *et al*., 2005), and ethyl parathion (Buratti *et al*., 2002).

In this work, we found that CYP2C9 was more selective towards thions compared to oxons using LC-MS/MS analysis. to. Nonetheless, the CWNA surrogate oxons, specifically CMP (cyclosarin surrogate) and PiMP (soman surrogate), were metabolized by CYP2C9. It was determined that the CYP2C9 inhibition potency changed with dose and time for nearly all OPs. We also performed *in silico* docking of the OP compounds crystal structure to investigate the correlation between metabolism, inhibition, and the computed binding energy (BE). This *in vitro* data could potentially be used to develop an artificial intelligence/machine learning-based (AI/ML) model for the prediction of metabolism of OP compounds by CYP450 enzymes. This is of particular importance as new OP threats are always emerging and will likely be aided by the use of AI/ML as demonstrated by the recent AI/ML-based prediction of 40,000 new OP structures (with many being more toxic) from the CWNA VX (Urbina *et al*., 2022, 2023).

## Materials and Methods

### Chemicals

All OP pesticides were purchased in neat form and used without purification. Phosfolan (CAS: 947-02-4) and tetrachlorvinphos (TCVP, CAS: 22248-79-9) were purchased from AccuStandard, Inc. (New Haven, CT, USA). Bensulide (BEN, CAS: 741-58-2), Chlorfenvinphos (CFVP, CAS: 470-90-6), Chlorphoxim (CPH, CAS: 14816-20-7), Chlorpyrifos Oxon (CPO, CAS: 5598-15-2), Crotoxyphos (CTP, CAS: 7700-17-6), Crufomate (CFA, CAS: 299-86-5), Cyanofenphos (CFP, CAS: 13067-93-1), Diazinon (DIA, CAS: 333-41-5), Dimefox (DME, CAS: 115-26-4), Formothion (FMN, CAS: 2540-82-1), Iodofenphos (CAS: 18181-70-9), Isofenphos (IFP, CAS: 25311-71-1), Isoxathion (IXT, CAS: 18854-01-8), Methidathion (MDT, CAS: 950-37-8), Mevinphos (MVP, CAS: 7786-34-7), Phosalone (PHO, CAS: 2310-17-0), Pyrazophos (PZP, CAS: 13457-18-6), Schradan (SCN, CAS: 152-16-9), Triazophos (TAP, CAS: 24017-47-8) and Tribufos (TBS, CAS: 78-48-8) were purchased from LGC Limited (Manchester, NH, USA). Acephate (ACE, CAS: 30560-19-1), Azinphos-Methyl (APM, CAS: 86-50-0), Chlorpyrifos (CPY, CAS: 2921-88-22), Dichlorvos (DCV, CAS: 62-73-7), Dimethoate (DMA, CAS: 60-51-5), Ethoprophos (EPP, CAS: 13194-48-4), Fenamiphos (FMP, CAS: 22224-92-6), Fenthion (FNN, CAS: 55-38-9), Iprobenfos (IBF, CAS: 26087-47-8), Malathion (MAL, CAS: 121-75-5), Mephosfolan (MPF, CAS: 950-10-7), Methamidophos (MMP, CAS: 10265-92-6), Monocrotophos (MCP, CAS: 6923-22-4), Phosmet (PMT, CAS: 732-11-6), Phosphamidon (PPM, CAS: 13171-21-6), Primiphos-ethyl (PPE, CAS: 23505-41-1), Pyridaphenthion (PPT, CAS: 119-12-0), Quinalphos (QUI, CAS: 13593-03-8), Temephos (TEM, CAS: 3383-96-8), and Trichlorfon (TCP, CAS: 52-68-6) were purchased from Sigma-Aldrich (Milwaukee, WI, USA). Sulfaphenazole (CAS: 526-08-9), NADPH (CAS: 53-59-8), and MgCl_2 (_CAS: 7791-18-6) were purchased from Sigma-Aldrich and were HPLC grade. The CWNA surrogates 3-cyano-4-methyl-2-oxo-2H-chromen-7-yl ethyl methylphosphonate (EMP, coumarin surrogate of VX), 3-cyano-4-methyl-2-oxo-2H-chromen-7-yl cyclohexyl methylphosphonate (CMP, coumarin surrogate of cyclosarin), 3-cyano-4-methyl-2-oxo-2H-chromen-7-yl (3,3-dimethylbutan-2-yl) methylphosphonate (PiMP, coumarin surrogate of soman), and ethyl (4-nitrophenyl) dimethylphosphoramidate (NEDPA, *p*-nitrophenol surrogate of tabun) were synthesized and generously supplied by Dr. Christopher Hadad and his research team at Ohio State University.

The OP structures used in this study are presented as oxons or thions in Figures S1 and S2 (see supporting information). All of the OP derivatives containing *E*/*Z* or *R*/*S* conformations were used as a mixture of isomers.

### pFluor50 CYP2C9 inhibition assay using CypExpress™

CypExpress™ 2C9 (Sigma, MTOXCE2C9), containing full length human recombinant CYP2C9, recombinant human P450 NADPH oxidoreductase, and a proprietary mix of MgCl_2_, glucose-6-phosphate dehydrogenase and glucose-6-phosphate, was purchased from Sigma Aldrich (St. Louis, MO). Magnesium chloride and β-nicotinamide adenine dinucleotide phosphate hydrate were purchased from Sigma Aldrich (St. Louis, MO) as described above. The fluorogenic substrate 7-ethoxy-3H-phenoxazin-3-one or resorufin ethoxy ether (Eres) (# 16122) was purchased from Cayman Chemical (Ann Arbor, Michigan). Perkin Elmer (Waltham, MA) OptiPlate-384 Black, Black Opaque 384-well Microplates (part no. 6007270) were used for all fluorogenic experiments. MilliQ water (18.2 MΩ) was used to prepare 0.1 M potassium phosphate buffer (pH 7.4) with potassium phosphate dibasic (0.06958 M) and potassium phosphate monobasic (0.03042 M). OP stocks were prepared at 0.5 mM (10 µM working concentration) in DMSO which were further diluted to 0.05 mM (1 µM final working concentration). Then, 1.2 µl (0.5 or 0.05 mM) of each OP stock was added to 28.8 µl of enzyme-containing master mix, comprised of 2 mg/ml CYP2C9, 0.65 mM NADPH, and 3.3 mM MgCl_2_. Compounds were pre-incubated with the master mix for 10 or 30 minutes. 30 µl of 4 µM Eres was then added to the master mix with OPs in each well after pre-incubation time was complete Fluorescent metabolite formation was determined by measuring the fluorescence (Ex 535, Em 590) on a BioTek Synergy H1 plate reader for 40 minutes under the following conditions: speed (normal); delay (100ms); measurements / read (100/read); read time interval (2 minute); height (9.75 mm) and gain (100). Sulfaphenazole was used as negative control for inhibition of CYP2C9. The slope was calculated as the change in relative fluorescent units over time, and data were normalized to the average slope of a vehicle control and % inhibition was calculated using Equation 1.

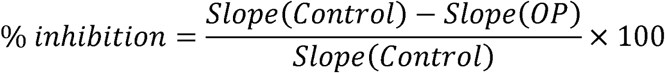

All data analyses were performed using Prism 10.4.2 software (GraphPad software Inc.). Experiments were performed with at least three technical replicates per experimental condition, and the reported values are the average of the multiple replicates. Pairwise-comparisons were performed using a Mann-Whitney test, with the Holm-Šídák method of multiple comparisons and an α value of 0.05 was used to determine significance.

### LC-MS/MS metabolism

*CYP2C9 Clearance:* CYP2C9 metabolism experiments were performed by BioDuro (Shanghai, China). Recombinant CYP2C9 (15 pmol) in phosphate buffer was pre-incubated with OPs (1 µM final reaction) at 37° C for 5 minutes before addition of NADPH (1 mM final reaction). Ten percent of the reaction was removed and quenched with a 5/10 ng/ml terfenadine/tolbutamide solution in acetonitrile at 0, 5, 30, 60 and 120 minutes. Samples were centrifuged at 4000 rpm at 4°C for 15 minutes and the supernatant was mixed 1:1 with water for LC-MS/MS analysis on either a Q Trap 4500 or API 4000. Diclofenac was used as the positive control CYP2C9 substrate. Parameters were calculated as follows:

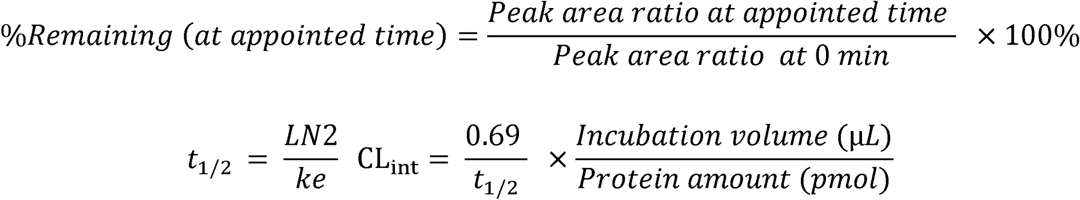

### Molecular docking analysis

#### Protein and Ligand Preparation

The crystal structure of human Cytochrome P450CYP2C9 with bound warfarin (PDB 1OG5) was used for subsequent molecular docking simulations (Williams *et al*., 2003). The structure was imported into the Molecular Operating Environment (MOE) software, where chain A was isolated and warfarin was removed (*Chemical Computing Group (CCG) | Research*, 2024).

Protein preparation was performed using MOE’s *QuickPrep* function, which applies homology modeling to complete missing residues, adds hydrogens, determines protonation states, and refines the structure using the Amber14:EHT force field (Labute, 2009; Maier *et al*., 2015). The receptor was then converted to pdbqt format using the utility tool *prepare_receptor* in the ADFR suite (Ravindranath *et al*., 2015). For the ligands, the OP pesticides’ SMILES strings were imported into MOE and prepared using the *Wash* method, where hydrogens were added, and the 3D coordinates were reconstructed. An energy minimization was performed with the Amber14:EHT forcefield (Maier *et al*., 2015). The prepared compound structures were then converted to pdbqt format using Python program Meeko, which supports atom typing and enhanced sampling of macrocycles during docking (Forli and Botta, 2007).

### Docking Method

Molecular docking calculations were conducted using Autodock Vina version 1.1.2, with the grid space centered at –22.89, 88.38, 33.046 Å and expanded 18, 20, and 20 points in the x, y, and z directions, respectively, to encompass the iron HEME and nearby residues. The protein was kept rigid during the docking calculations. Before docking the OP pesticides, warfarin was re-docked onto its crystallographic binding site to validate the protocol. The top scored pose had the best alignment with the experimental conformation (Williams *et al*., 2003). Subsequently, each OP pesticide was docked into the receptor at an exhaustive level of 200, generating up to 20 unique poses.

### Correlation analysis

Time-dependent differences in inhibition were calculated from the 1 µM and 10 µM inhibition experiments with preincubation of the substrate for either 10 or 30 minutes. Similarly, the percentage of parent remaining after 2 hours in the LC-MS/MS was determined. Correlation analyses were performed in Graphpad Prism v10.4.2. We hypothesized that the iron center would interact with the double bonded oxygen or sulfur, so from the docking calculations, a reactive distance was computed in order to represent the distance between the P=O for oxons or P=S for thions to the iron atom in the HEME (i.e. the reactive distance) for all poses and then the binding energies of those poses were compiled. For each OP, the BE of the pose with the shortest reactive distance was used in all subsequent analysis. Binding efficiency (BE_eff_), the BE divided by the heavy atom molecular weight, and the shortest distance were correlated against the time-dependent difference in inhibition or percentage of parent remaining. For each correlation analysis, the R-value was obtained from the equation for the linear fit of the data.

## Results

### CYP2C9 activity is inhibited by thions and oxons

We used our previously developed pFluor50 high-throughput fluorogenic assay to determine inhibition of CYP2C9 activity by OP compounds using Eres as a substrate and sulfaphenazole as a positive control inhibitor (Shriwas et. al. 2025). We performed the activity assay with 47 different OP derivatives to determine if they are inhibitors or activators of CYP2C9.

Sulfaphenzole was used as positive control as it almost completely inhibits CYP2C9 activity at the concentrations used (Figure 1A, B). We determined that out of 21 thions, 18 at 10 µM and 12 at 1 µM exhibited significant time-dependent variations in inhibition when preincubated for 10 or 30 minutes (Figure 1 A, B). While out of 26 oxons, 23 at 10 µM and 10 at 1 µM exhibited significant time-dependent variations in inhibition between 10 and 30 minutes, respectively (Figure 1 A, B). The data are also summarized in a heatmap of the OP-mediated inhibition of CYP2C9 activity (Figure 1C). Overall, we found that several OP pesticides (CPY, PHO, EPP, IBF, MPF, etc.) were activators of CYP2C9 after 10-minutes of preincubation but became inhibitors after 30-minutes of preincubation. Further, some OP pesticides, such as BEN and CPO, showed a time-dependent decrease in inhibition. Time-dependent increases in inhibition suggest either mechanism-based OP inhibition of CYP2C9 with a slow second step or more likely that the OP is metabolized by CYP2C9 leading to a product/metabolite that is more inhibitory.

**Figure 1.**
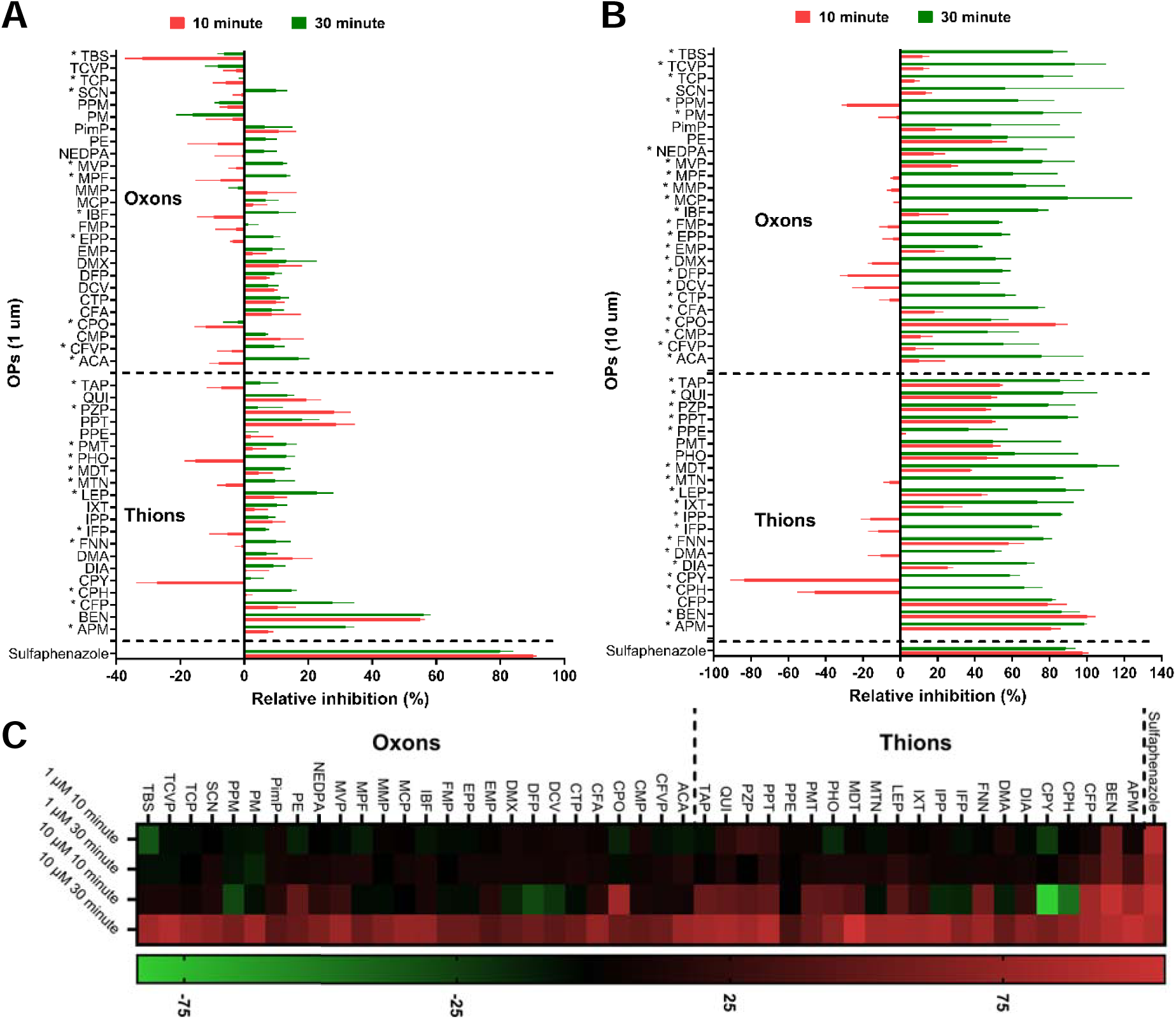
OP-mediated relative inhibition using pflour50 fluorogenic assay for CYP2C9 activity. **A**. Relative inhibition of CYP2C9 activity by thions and oxons at 1 µM concentration. **B**. Relative inhibition of CYP2C9 activity by thions and oxons at 10 µM concentration. **C**. Heatmap showing variations in CYP2C9 activity due to OP inhibition at 1 and 10 µM between 10 and 30 min. * Represents OP compounds wherein the inhibition of CYP2C9 changes significantly between 10 and 30 min. Sulfaphenazole was used as positive control for inhibition of CYP2C9.

Decreases in inhibition suggest metabolism with the product/metabolite being less inhibitory toward CYP2C9.

### Thions are selectively metabolized by CYP2C9 (as compared to oxons)

We analyzed the metabolism of thions and oxons (including CWNA surrogates) as determined by the percentage of parent OP remaining from LC-MS/MS experiments and determined the intrinsic clearance rate. We observed that all 12 thions studied were significantly degraded by CYP2C9 after 2 hours with MDT being metabolized the slowest but still degraded by at least 35% (Figure 2B). FMN was completely degraded within 30 minutes while DIA, PHO, IXT, QPS and TEM were metabolized by ∼80% after 2 hours (Figure 2B). Among the 10 OP pesticides as oxons, only FMP was metabolized completely (Figure 2A), whereas TCP, TCVP, CFVP and EPP were metabolized by ∼35%. The other OP pesticide oxons which were studied were not significantly metabolized within the time studied. The tested CWNA surrogates as OP oxons were only incubated with CYP2C9 for 1 hour (Figure 2C); however, CMP (cyclosarin surrogate) and PiMP (soman surrogate) were metabolized by ≥ 35%. The intrinsic clearance rate calculated for each of the OP compounds are listed in Table 1.

**Figure 2.**
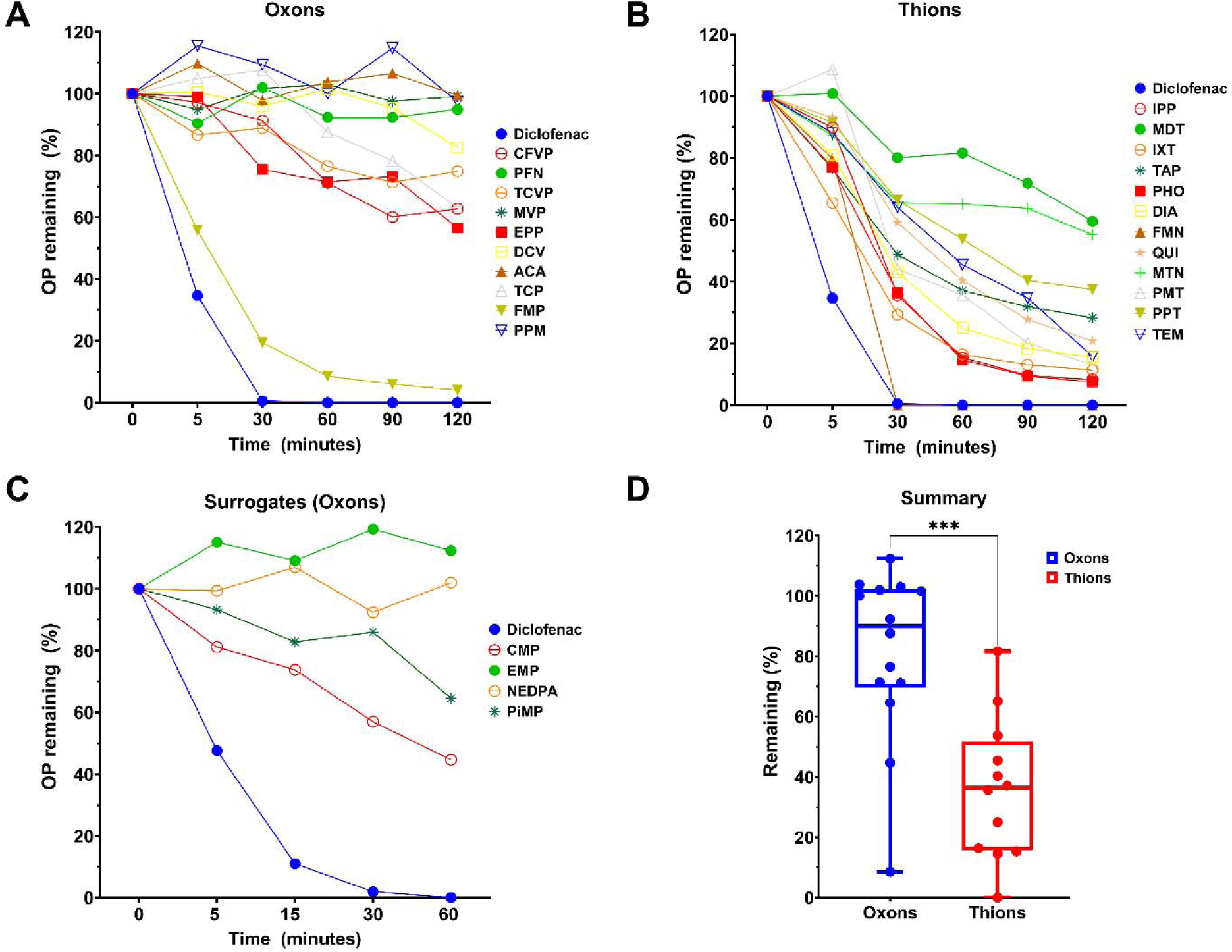
Metabolism of OP compounds by CYP2C9 measured by disappearance of parent. **A**. LC-MS/MS degradation of oxons by CYP2C9 after 2 hrs. **B**. LC-MS/MS degradation of thions by CYP2C9 after 2 hrs. **C**. LC-MS/MS degradation of CWNA surrogates after 1 hour. **D.** CYP2C9 mediated degradation of thions was significantly more as compared to oxons based on mean averages after 2 hrs.

**Table 1.** Half-life and intrinsic clearance of OP pesticides by CYP2C19 as measured by disappearance of parent.

| Compound | $t_{1/2}$ (min) | $CL_{int}$ (μL/min /pmol CYP) |
| --- | --- | --- |
| Diclofenac | <5 | >2.8 |
| **Oxon** |  |  |
| PFN, DCV, ACA, PPM, MVP | >372.8 | <0.04 |
| EMP, NEDPA | >186.4 | <0.07 |
| CFVP | 153.3 | 0.1 |
| TCVP | 214.3 | 0.4 |
| TCP | 172.2 | 0.1 |
| EPP | 164.9 | 0.1 |
| CMP | 54.85 | 0.25 |
| PiMP | 105.46 | 0.13 |
| FMP | 27.1 | 0.5 |
| **Thion** |  |  |
| IPP | 31.3 | 0.4 |
| MDT | 171.6 | 0.1 |
| PHO | 31.2 | 0.4 |
| IXT | 31.5 | 0.4 |
| TAP | 57.3 | 0.2 |
| DIA | 44.2 | 0.3 |
| FON | 15.3 | 0.9 |
| PHT | 39.8 | 0.3 |
| PPT | 82.6 | 0.2 |
| TEM | 49.4 | 0.3 |
| QPS | 52.2 | 0.3 |
| MTN | 52.4 | 0.3 |

### Molecular docking analysis reveals correlation between binding energy and degradation or time-dependent change in inhibition

We performed molecular docking using Autodock Vina with the OP compounds studied experimentally (Figures S1-S2). Docking was performed with CYP2C9 (PDB 1OG5) which was selected based on the resolution of the structure and the bound ligand. We then performed a correlation analysis of the percentage of parent OP remaining after 2 hours with the computed BE from the pose with the shortest reactive distance. We also explored the correlation of the experimental data with the shortest reactive distance between the OP atom and the iron atom in the heme ring. Additionally, a binding efficiency (BE_eff_) was computed representing the BE divided by the OP’s heavy atom molecular weight. Notably, we identified that the computed BE of the pose with the shortest reactive distance yielded the best correlation to the experimental data. Figure 3A shows the correlation between the percent of parent remaining and the BE for the pose with the shortest reactive distance with a correlation coefficient of R 0.61. This analysis excluded CWNA surrogates as their metabolism was only tested for 1 hour. The OP compounds which were metabolized to a greater extent had lower binding energies while the OP compounds which were not metabolized had higher binding energies. Thus, OP compouns could be clustered into two groups (Figure 3A) except for FMN, PPT, CFVP and TCVP. We also performed a similar analysis with the degradation data at 1 hour (which included the surrogates) but found a much lower correlation (R0.38) (Figure S3A). The correlation between the degradation and the shortest reactive distance and the binding efficiency for both 2 hour and 1 hour data can be seen in Figure S3B-E. Next, we performed a correlation analysis between the time-dependent change in inhibition (between 10 and 30 minutes of preincubation) at the 10 µM OP concentration with the computed binding energy of the pose with the shortest reactive distance. We again obtained positive correlation with correlation coefficient of R0.59 (Figure 3B). Unlike LC-MS analysis, which was performed for 22 OP compounds, this analysis included all 47 OP derivatives. Two notable outliers were removed from this analysis: IPP and CPY had the greatest time-dependent changes in inhibition between the 10-and 30-minute preincubation. When we analyzed the correlation between the time-dependent change in inhibition with the calculated binding energy for just the oxons, we obtained R = 0.56 (Figure S4A), whereas a similar analysis with only the thions resulted in R0.63 (after exclusion of the previously noted outliers; Figure S4B). Similar analysis was performed for the 1 µM OP time-dependent change in inhibition and the calculated binding energy (Figure S3C). TAP was the OP that demonstrated the strongest correlation with the *in silico* binding energy compared to the LC-MS/MS degradation and inhibition potential (Figure 3C-D). Similarly, we performed a correlation analysis for the time-dependent change in inhibition at the 10 µM OP concentration with the shortest reactive distance between the OP and the heme ring center and the binding efficiency (Figure S3D-E). Overall, the R values of 0.59, 0.56 and 0.63 showed that the docking studies may have value in predicting OP metabolism.

**Figure 3.**
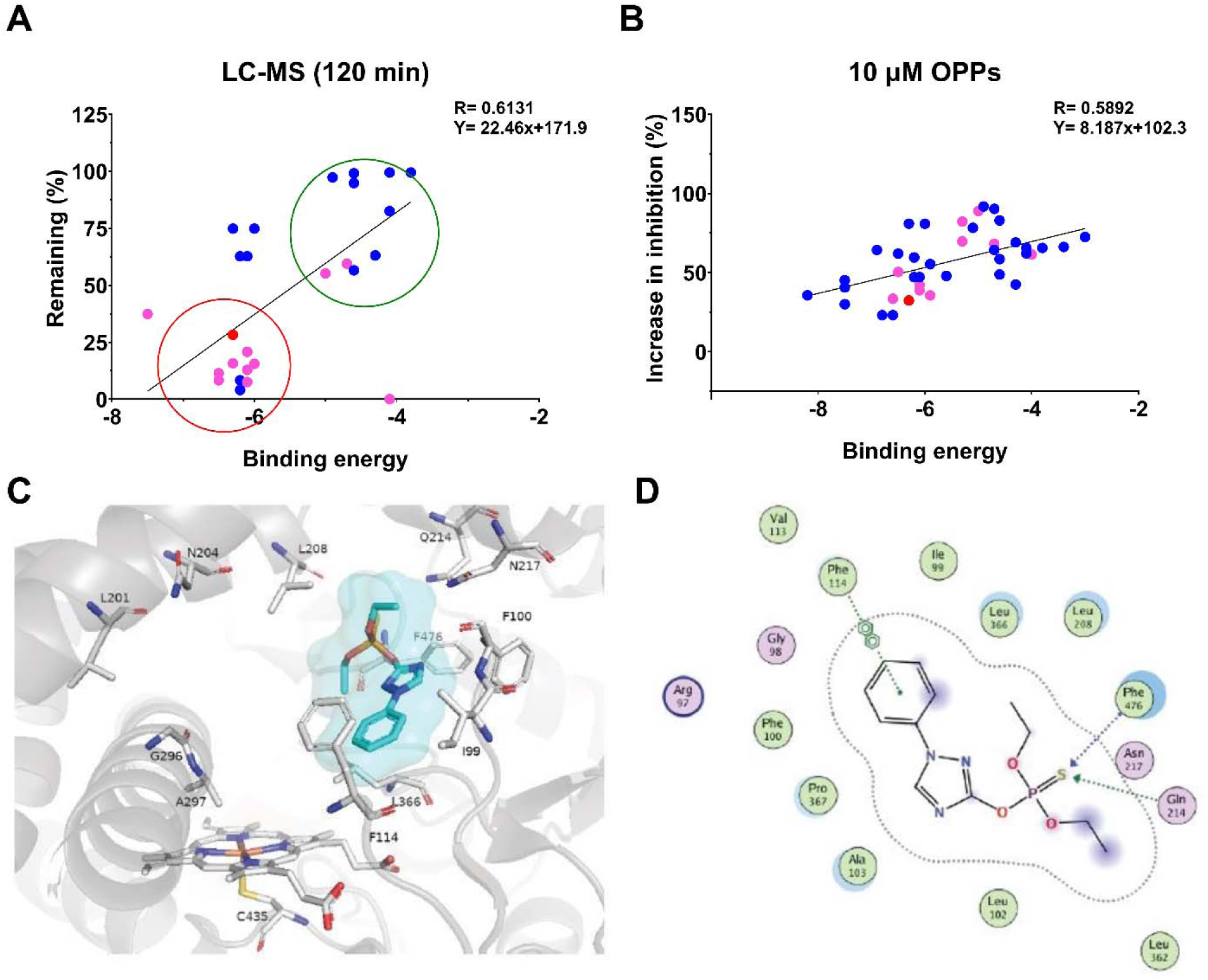
**Correlation analysis between binding energy (docking) vs experimental results**. **A**. Correlation analysis (R=0.6131; P<0.0005) between percent of parent remaining in the LC-MS/MS metabolism experiments after 2 hrs and the binding energy in the docking experiments. Pink circles represent thions, blue circles oxons, and the red circle is TAP. **B**. Correlation analysis (R= 0.5892; P<0.0001) between time-dependent increase in inhibition (%) in the pFlour50 assay and the binding energy. Pink circles represent thions, blue circles oxons, and the red circle is TAP. **C.** TAP docking into 1OG5 active site nearly perpendicular to heme ring. **D.** 2-D interaction diagram of TAP with 1OG5 amino acids showing Phe114 (П-П) and Phe476/Gln214 interaction with the sulfur.

## Discussion

Organophosphate (OP) poisoning is among the leading causes of suicide worldwide, accounting for approximately 15% of all suicide-related deaths globally (Mew *et al*., 2017). When accidental acute OP poisoning is also considered, the annual death toll rises to an estimated 150,000– 200,000 cases (Blain, 2011; Eddleston, 2024). Beyond fatalities, the widespread use of OP pesticides in agriculture exposes workers frequently and results in residues on produce such as fruits and vegetables, contributing to chronic OP exposure in a large portion of the population (Beyuo *et al*., 2024; Habran *et al*., 2024; Soman *et al*., 2024). Existing treatments for OP poisoning remain insufficient, especially regarding effects on the central nervous system. As a result, gaining a deeper understanding of how OP compounds are metabolized is crucial.

We previously demonstrated that human liver microsomes (HLMs) could metabolize many OP compounds using targeted LC-MS/MS and NMR-based metabolomics (Agarwal *et al*., 2023). However, HLMs contain many metabolizing enzymes including CYP450s which are naturally occurring bioscavenging enzymes that primarily regulate xenobiotic drug metabolism in the human body (Zhao *et al*., 2021). We have now studied the metabolism of OP compounds by CYP2C9 using LC-MS/MS to determine degradation of 26 OP derivatives including 4 CWNA surrogates.

We found that thions were more readily metabolized as compared to oxons (Figure 2D). Thions are generally converted to the active metabolite oxon through oxidation of the phosphorus-bound sulfur to the corresponding oxygen. CYP2C9 has been previously been shown to desulfurate thions, such as diazinon (Kappers *et al*., 2001), dimethoate (Buratti and Testai, 2007), disulfoton (Usmani *et al*., 2004), fenthion (Leoni, Buratti and Testai, 2008), malathion (Buratti *et al*., 2005), and ethyl parathion (Buratti *et al*., 2002). Here we extend these findings to additional OP pesticides while also studying the metabolism of oxons. Particularly, our work confirms and extends on the previously demonstrated metabolism of diazinon and malathion by CYP2C9 (Figure 2B). Correlation analysis between the percent of parent OP remaining after 2 hours with the computed binding energy (BE) showed a correlation coefficient of R0.61 suggesting tighter binding leads to greater degradation. CFVP, TCVP and formothion were outliers, which could be likely due to formation of secondary metabolites that inhibit further degradation by CYP2C9. On average, the BE for thions showed more favorable binding as compared to the oxons. However, the BE (–5.7 kcal/mol) of the thion CPY and its corresponding oxon CPO, were very similar, suggesting that moieties other than just the sulfur versus oxygen on the phosphorus of the OP compound may contribute to the interaction with CYP2C9. Based on the *in vitro* data and our docking insights, the correlations suggest that we may be able to develop a model based on OP structure to predict CYP2C9 metabolism of new OP compounds.

We previously developed a fluorogenic assay for CYP2C9 using ethoxyresorufin as a substrate, pFlour50 (Shriwas *et al*., 2025). We further showed that sulfaphenazole inhibits CYP2C9 in a dose dependent, but time-independent manner. Using this previously developed method, we determined the time-dependent CYP2C9 activation and/or inhibition by OP compounds. We performed the inhibition studies at two different concentrations (1 and 10 µM) and two different preincubation times (10 and 30 minutes) to study the time and dose dependence of the inhibition. We found that at the 10 µM OP concentration, there was a significant change in CYP2C9 inhibition by a large proportion of the OP compounds studied, indicating either mechanism-based inhibition or metabolism by CYP2C9 to secondary metabolites with stronger binding affinity. Furthermore, these changes were not as prevalent at the 1 µM OP concentration, suggesting that the affinity of CYP2C9 for the OP compounds may be rather low.

Docking analysis revealed that the time-dependent change in inhibition was positively correlated with the BE (R= 0.59) of the pose with the shortest reactive distance. The majority of the oxons studied fit well to the correlation curve except for PE and CPO. PE showed almost no time-dependent change in inhibition whereas CPO showed a decrease in inhibition with time, suggesting that it is metabolized to a product with less inhibition of CYP2C9. Interestingly, CPY (the corresponding thion of the oxon CPO) inhibition after the 10-minute preincubation demonstrated it was an activator (∼30%) whereas after the 30-minute preincubation was ∼59%.

CPO inhibition at the same time was ∼83%. This suggests that CPY becomes metabolized after 30 minutes to a mix of CPY and CPO resulting in greater inhibition of CYP2C9. For the thions studied, there were multiple outliers such as PHO, PMT and IPP which also demonstrated significant metabolism in the LC-MS/MS studies, suggesting that these compounds are metabolized into secondary metabolites that are more inhibitory towards CYP2C9. Similarly, this might be the reason for the variations we observed with other thions such as APM, BEN, CPH, CPY, CFP and fenthion. Overall, the positive correlation observed (R = 0.61) indicates that it might be possible to predict the inhibitory activity based on the computed BE. Repeating these experiments with additional OP compounds might improve the predictive power in the future.

The OPNA surrogate data analysis demonstrated that CMP (a surrogate of cyclosarin) and PiMP (a soman surrogate) were both metabolized by CYP2C9 after 1 hour. Furthermore, CMP had the largest van der Waal radii, which may have contributed to stronger interactions leading to its BE, which was the lowest of those studied. Thus, comparing the binding sites of CMP versus cyclosarin, PiMP versus soman, and NEDPA versus tabun showed multiple differences suggesting that these surrogates might not represent the CWNAs well and further that CYP2C9 may not metabolize the authentic CWNAs. However, VX and EMP (a surrogate of VX) bound at the same site and had very similar interactions with the enzyme suggesting that EMP is likely predictive of VX, indicating that VX is likely not metabolized by CYP2C9 (since EMP was not).

## Conclusions

In conclusion, although CYP2C9 is likely responsible for the metabolism of the majority of thion OP pesticides, it likely is not involved in the metabolism of CWNAs. In this study, we were able to get moderate correlation but with more number of molecules to study we might get a much stronger correlation. Further, we have only performed docking on OPs but we could improve using more robust molecular dynamics simulation for degraded OPs. Recently, an AI/ML model was used to predict 40,000 new OP structures from VX, many of which were predicted to be more toxic than the starting structure (Urbina *et al*., 2022, 2023). While experimental analysis of all of these theoretical structures is not possible, the experiments and computational analysis contained in the current study suggest that through analysis of the susceptibility of a subset of OP compounds to degradation by and time-dependent changes in inhibition of the CYP450s along with the predicted BE it may be possible to develop AI/ML-based models to predict the metabolism of the OP compounds. In the future, we will explore similar analyses of OPs with other CYPs to determine if these results are specific to CYP2C9 or are more broadly applicable to all of the relevant human CYP450s.

## Abbrevations

AChE: Acetylcholinesterase
ACA: Acephate
APM: Azinphos-methyl
BE: binding energy
BNS: Bensulide
CFVP: Chlorfenvinphos
CPH: Chlorphoxim
CPY: Chlorpyrifos
CPO: Chlorpyrifos Oxon
CTP: Crotoxyphos
CFA: Crufomate
CFP: Cyanofenphos
CYP: cytochrome p450
CWNA: chemical warfare nerve agent
DFP: Diisopropyl fluorophosphate
DIZ: Diazinon
DCV: Dichlorvos
DEP: Diethylparaoxon
DFP: Diisopropyl fluorophosphate
DMX: Dimefox
DMA: Dimethoate
DMP: Dimethyl paraoxon
EPP: Ethoprophos
Eres: resorufin ethoxy ether
FMP: Fenamiphos
FNN: Fenthion
FON: Formothion
IFP: Iodofenphos
IBF: Iprobenfos
IPP: Isofenphos
IXT: Isoxathion
LPS: Leptophos
MTN: Malathion
MPF: Mephosfolan
MMP: Methamidophos
MDT: Methidathion
MVP: Mevinphos
MCP: Monocrotophos
OP: organophosphorus compound
PHO: Phosalone
PFN: Phosfolan
PHT: Phosmet
PPM: Phosphamidon
PPE: Pirimiphos-ethyl
PZP: Pyrazophos
PPT: Pyridaphenthion
QNP: Quinalphos
SCN: Schradan
Sul: Sulfaphenazole
TEM: Temephos
TCVP: Tetrachlorvinphos
TAP: Triazophos
TBS: Tribufos
TCP: Trichlorphon

## Patents

Not applicable.

## Supplementary Materials

Supplementary materials are attached separately.

## Author Contributions

Conceptualization, P.S. and C.M.; methodology, P.S. A.N.; software, P.S., A.N., and C.H.; validation, C.M., C.H., T.M. and S.E.; formal analysis, P.S. and C.M.; investigation, P.S., A.R., and A.N.; resources, C.M., C.H. and S.E.; data curation, P.S.; writing— original draft preparation, P.S.; writing—review and editing, P.S., C.M., C.H., T.L. and S.E; visualization, P.S.; supervision, C.M., C.H., and S.E.; project administration, C.M. and S.E..; funding acquisition, C.M. and S.E. All authors have read and agreed to the published version of the manuscript.

## Institutional Review Board Statement

Not applicable.

## Informed Consent Statement

Not applicable.

## Data Availability Statement

Data are contained within the article.

## Acknowledgments

The authors would like to thank Dmitry Uchenik (Shared Instrumentation Facility Director, College of Pharmacy) instrument facility for use of the BioTek plate reader.

