## Supplementary for "Human CYP2C9 metabolism of organophosphorus pesticides and nerve agent surrogates"

**Supplementary materials**

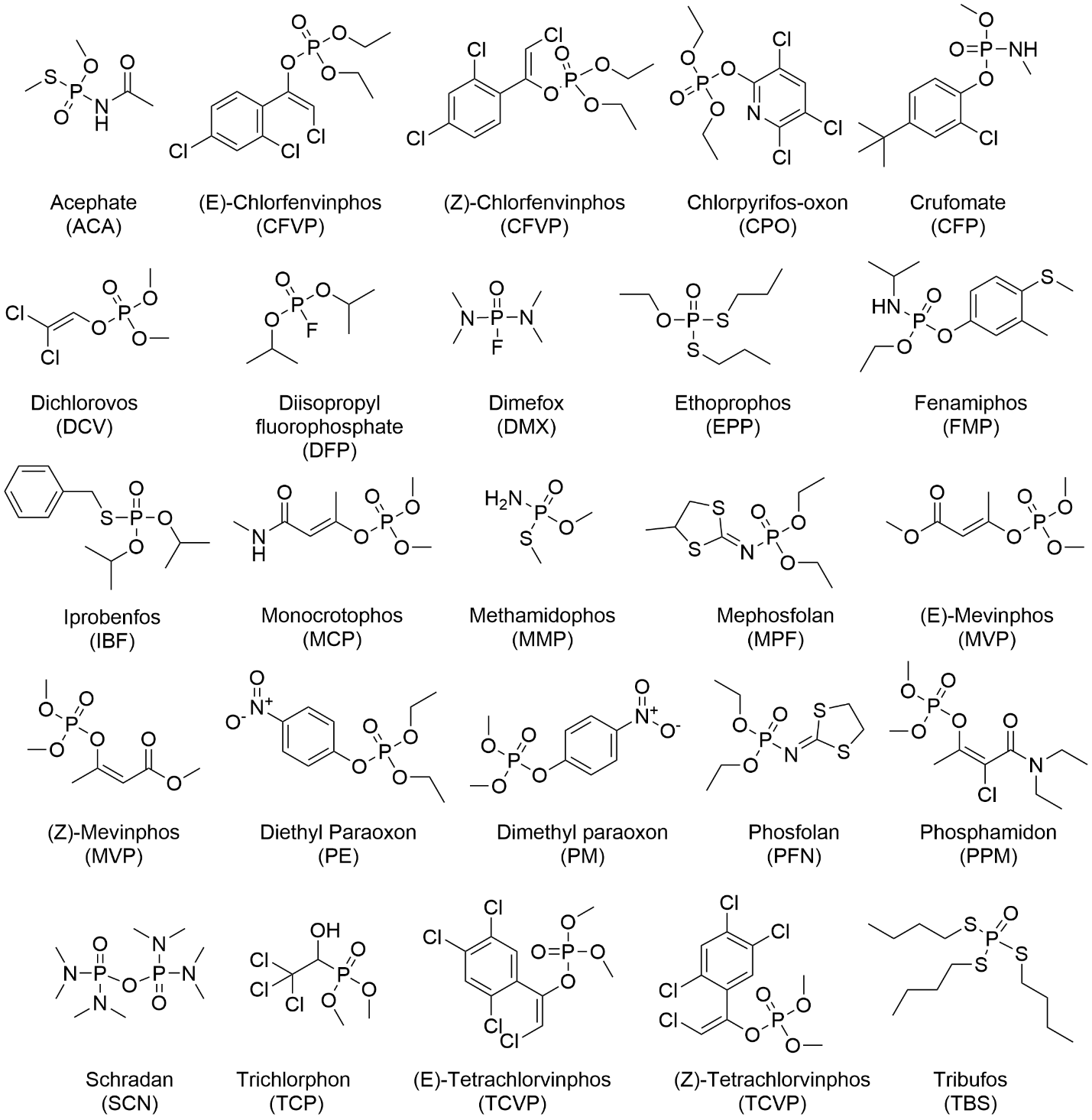

**Figure S1: Structures of the oxons used in experiments and docking analysis.**

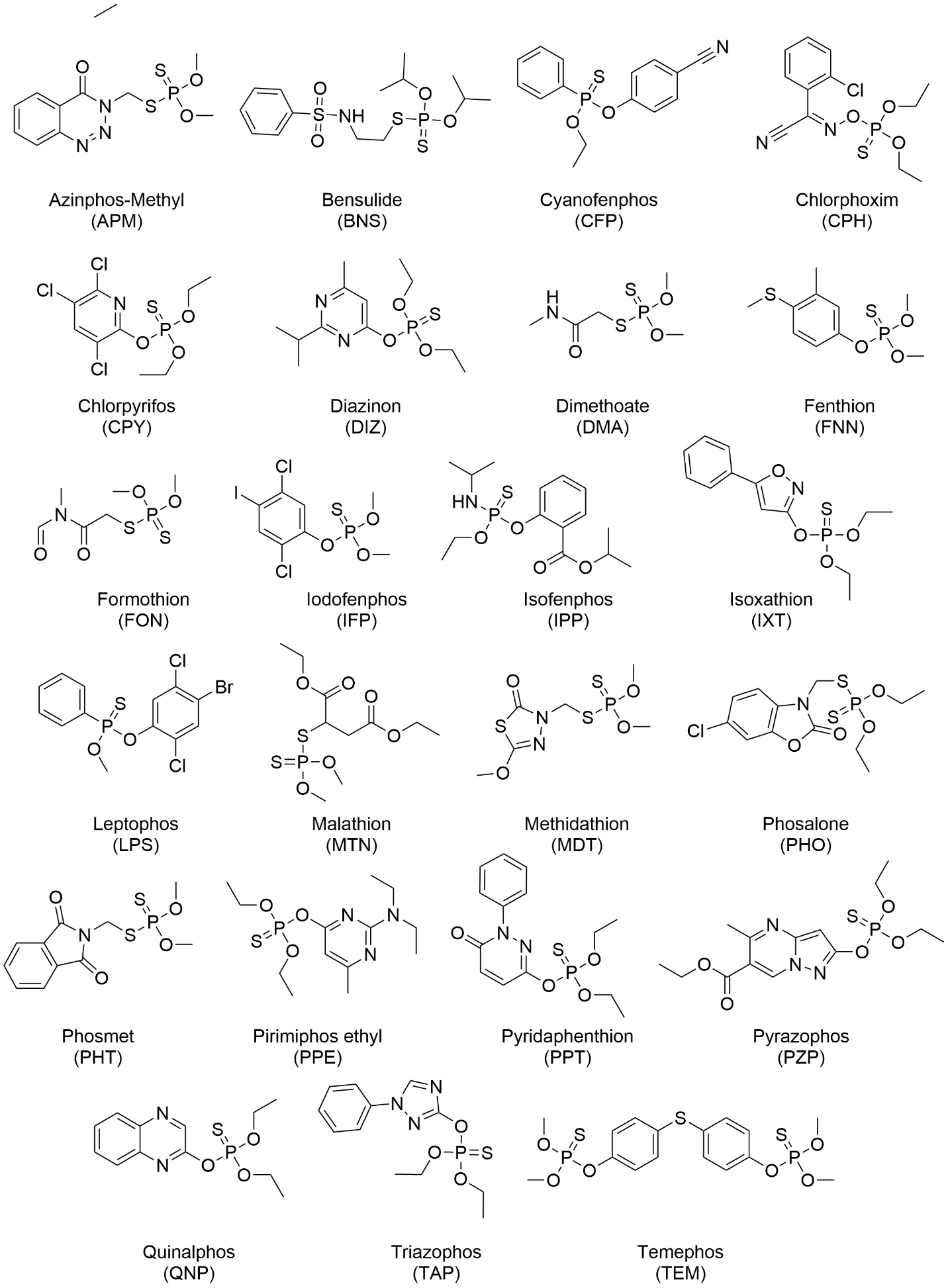

**Figure S2: Structures of the thions used in experiments and docking analysis**

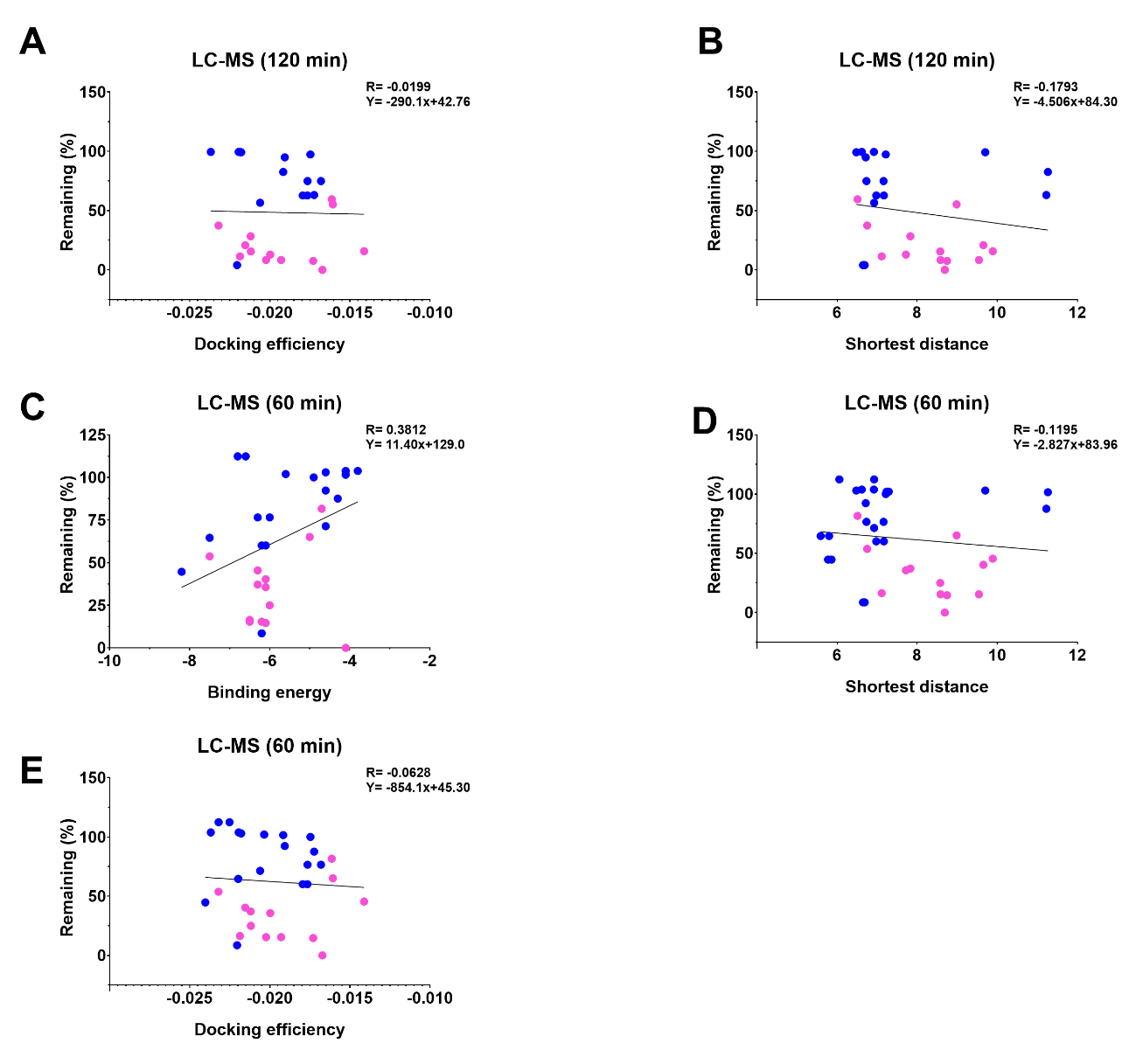

**Figure S3. Correlation between LC-MS/MS data with docking parameters**. **A**. Correlation analysis between metabolite remaining in LC-MS/MS metabolism (2 hours) and docking efficiency. **B**. Correlation analysis between metabolite remaining in LC-MS/MS metabolism (2 hours) and shortest distance. **C**. Correlation analysis between metabolite remaining in LC-MS/MS metabolism (1 hour) and binding energy. **D**. Correlation analysis between metabolite remaining in LC-MS/MS metabolism (1 hour) and shortest distance. **E**. Correlation analysis between metabolite remaining in LC-MS/MS metabolism (1 hour) and docking efficiency. Diclofenac was used as a control for CYP2C9 metabolism. Dot color: Pink: thion; Blue: oxon

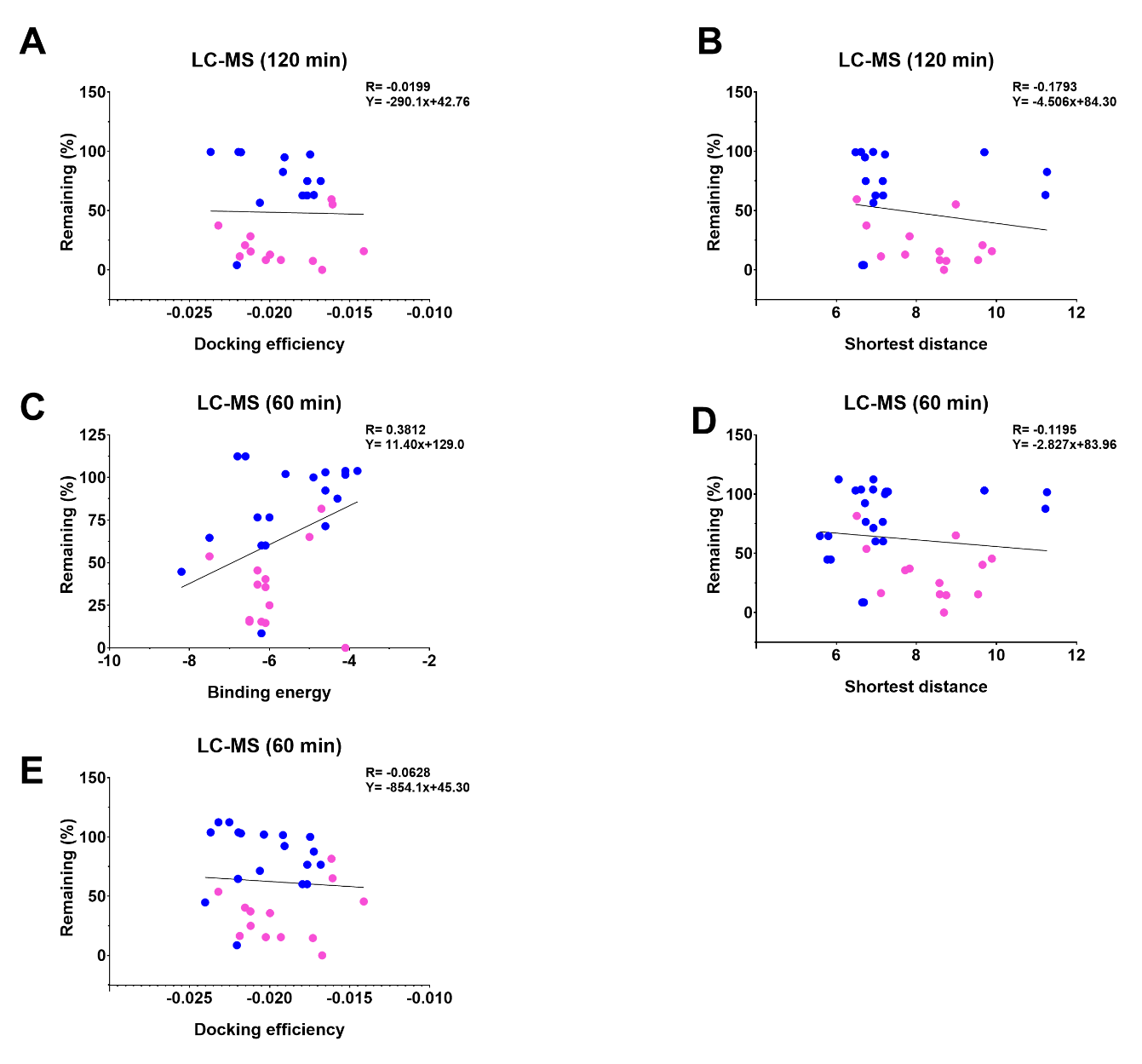

**Figure S4. Correlation between inhibition data with docking parameters**. **A**. Correlation analysis between inhibition data and binding energy for oxons at 10 µM concentration. **B**. Correlation analysis between inhibition data and binding energy for thions at 10 µM concentration. **C**. Correlation analysis between inhibition data and binding energy at 1 µM OP concentration. **D**. Correlation analysis between inhibition data and docking efficiency at 1 µM concentration. **E**. Correlation analysis between inhibition data and shortest distance at 1 µM concentration. Sulfaphenazole was used as a control for CYP2C9 metabolism. Dot color: Pink: thion; Blue: oxon
